# A Comparison of Machine Learning and Bayesian Modelling for Molecular Serotyping

**DOI:** 10.1101/138636

**Authors:** Richard Newton, Lorenz Wernisch

## Abstract

**Background:** *Streptococcus pneumoniae* is a human pathogen that is a major cause of infant mortality. Identifying the pneumococcal serotype is an important step in monitoring the impact of vaccines used to protect against disease. Genomic microarrays provide an effective method for molecular serotyping. Previously we developed an empirical Bayesian model for the classification of serotypes from a molecular serotyping array. With only few samples available, a model driven approach was the only option. In the meanwhile, several thousand samples have been made available to us, providing an opportunity to investigate serotype classification by machine learning methods, which could complement the Bayesian model.

**Results:** We compare the performance of the original Bayesian model with two machine learning algorithms: Gradient Boosting Machines and Random Forests. We present our results as an example of a generic strategy whereby a preliminary probabilistic model is complemented or replaced by a machine learning classifier once enough data are available. Despite the availability of thousands of serotyping arrays, a problem encountered when applying machine learning methods is the lack of training data containing mixtures of serotypes; due to the large number of possible combinations. Most of the available training data comprises samples with only a single serotype. To overcome the lack of training data we implemented an iterative analysis, creating artificial training data of serotype mixtures by combining raw data from single serotype arrays.

**Conclusions:** With the enhanced training set the machine learning algorithms out perform the original Bayesian model. However, for serotypes currently lacking sufficient training data the best performing implementation was a combination of the results of the Bayesian Model and the Gradient Boosting Machine. As well as being an effective method for classifying biological data, machine learning can also be used as an efficient method for revealing subtle biological insights, which we illustrate with an example.

## Background

We investigate different approaches to analysing the raw data from a custom genomic microarray that has been designed for molecular serotyping *Streptococcus pneumoniae* [1, 2]. *Streptococcus pneumoniae* is an important human pathogen and a major cause of infant mortality. There are over ninety known serotypes of the bacterium and there is a requirement to monitor the population dynamics of the different *S. pneumoniae* serotypes worldwide. A custom molecular serotyping array [3] was developed to fulfil the need for an accurate and objective serotyping method which could easily detect multiple serotypes in a sample. The method gave the best performance of any serotyping method in the independent Pneucarriage Project [2]. Genomic microarrays are a method for detecting the presence or absence of multiple genes within a sample simultaneously, through specific binding to an array of high-density probes. A microarray constructed with probes for genes specific to different strains of an organism can detect the presence of a particular strain of the organism in a clinical sample according to which of the probes have an elevated signal.

The custom B*µ*GS SP-CPS molecular serotyping microarray [3] contains probes for over four hundred *S. pneumoniae* capsular polysaccharide synthesis (*cps*) genes. Each serotype is known to contain a small subset of these *cps* genes, ranging from 1 to 22 genes. However, there is considerable overlap between the subsets with no serotype having a set of genes unique to itself. In fact a few serotypes have identical, or near identical, subsets of *cps* genes, so in order to distinguish these serotypes the array contains further discriminative probes. An analysis method to produce a call as to which serotypes are present in a sample has to process these two types of probe and also allow for cross-hybridisation to probes and cope with the inevitable experimental noise. An empirical Bayesian model was an obvious choice of method, given this need to integrate data types with prior knowledge.

The model we developed [1] has proved to be very effective given the limited amount of sample data available at the time. There is however far more sample data now available, which has made possible the application of machine learning classifiers. The Bayesian modelling approach requires decisions on what prior knowledge should be included in the model; prior knowledge which may be subjective and incomplete. The main advantage of machine learning methods over a Bayesian model is that the important discriminative features are learnt automatically by the algorithm from a, usually large, number of training samples. The discriminative features that the algorithm learns may also reveal interesting biological insights previously hidden in the data. The main problems that may be encountered in a machine learning approach is insufficient training data to capture all the idiosyncrasies that may be encountered in test data, and overfitting of the training data.

There are now a considerable number of training samples available that contain single serotypes. However, an important requirement for any *Streptococcus pneumoniae* serotyping method is the ability to detect multiple serotypes in a sample. The main difficulty of applying machine learning to this particular problem is insufficient samples available containing all possible combinations of mixtures of serotypes. We overcame this limitation by implementing an two-step iterative approach which involves the construction of artificial training data.

The two machine learning methods investigated here are Random Forests (RF) and Gradient Boosting Machines (GBM). Random Forests is an ensemble learning method that uses decision trees to classify the data. To mitigate the tendency to overfit, many decision trees are fitted to the data and the mode of the predicted classes taken. Gradient Boosting Machines (GBMs) is also an ensemble learning method and in the current application, also uses decision trees. Unlike Random Forests, GBMs use boosting, that is, the ensemble of trees are fit in sequence and at each step the tree is fitted not to the data but to the residual error from the fit of the ensemble so far.

The purpose of the study was to evaluate the analysis of the molecular serotyping microarray by machine learning methods, firstly as a classifier but also as a biological research tool. A further purpose of this study is to propose some generic strategies for the application of machine learning methods, for example, how to combine a statistical model with a machine learning classifier, or how to compensate for the lack of training data due to a combinatorial explosion of classification options.

## Methods

### *Streptococcus pneumoniae* molecular serotyping microarray

The custom genomic microarray [3] contains several thousand oligonucleotide probes designed to detect a number of different entities (also see [1] for more details):

1. Probes, referred to here as CPS probes, for 441 capsular polysaccharide synthesis *cps* genes. On average there are 10 probes per gene. These probes are the primary source of information on the serotypes in the sample. Figure 1 shows schematically the relationship between a serotype, *cps* genes and the CPS probes on the array.
2. Probes, referred to here as STID probes, designed to identify serotypes that are too closely related to be resolved by the *cps* genes alone.
3. Probes on the microarray for the *entire* genome of *Streptococcus pneumoniae* from two sequenced strains of the bacterium, 6824 probes in total and referred to here as the *genome* probes. In the context of the current study these probes are used only to provide a reference level with which to normalise the arrays for the machine learning algorithms and in the derivation of priors for the Bayesian model.

**Figure 1.**
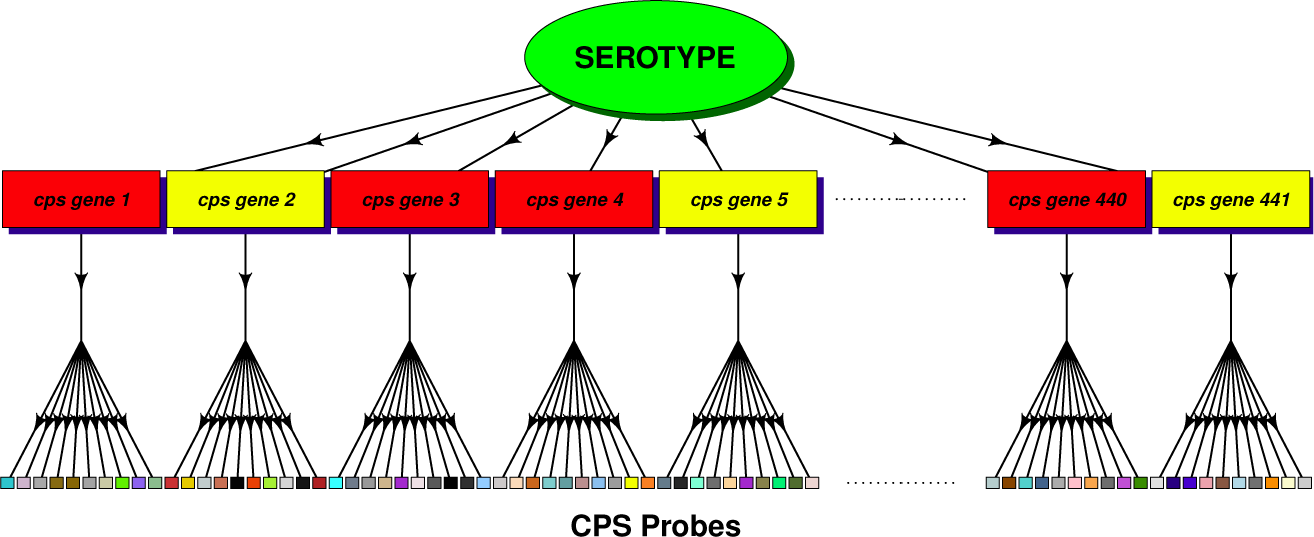
Diagram of the data structure. Schematic showing the relationship between a serotype, *cps* genes (those present in the serotype, coloured red, those absent coloured yellow) and the CPS probes on the array.

Figure 2 shows CPS probe data from a typical microarray testing a sample containing one serotype (17F). Figure 2a displays the raw probe intensities. In Figure 2b the probes have been grouped according to their target genes and displayed as a boxplot of log intensities. Only the top 75 genes, out of a total of 441 genes, with the highest median probe log intensity are plotted for clarity. In Figure 2c a t-test has been performed for the probes of each gene; testing whether the mean of the probes is significantly greater than the mean of all the other probes. The figure displays the *p*-values from the t-tests for the top 75 genes.

**Figure 2.**
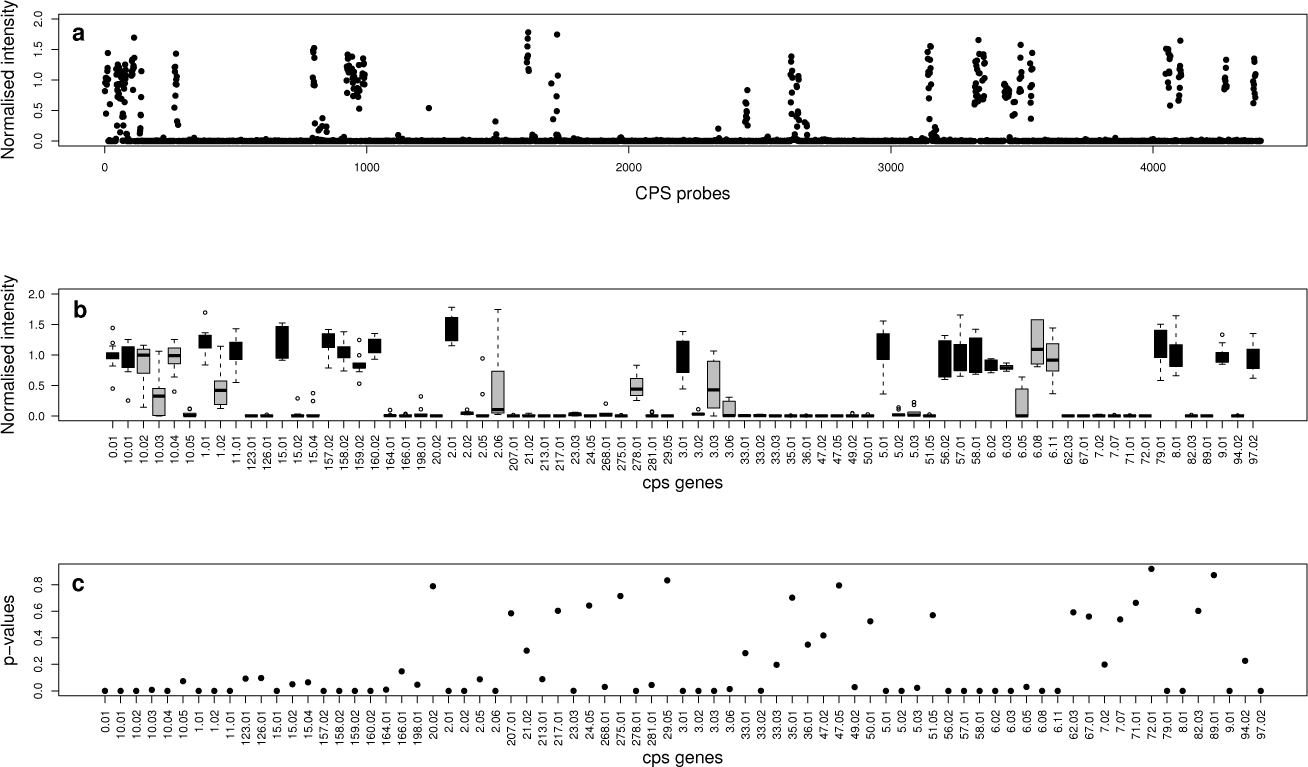
Array data from typical single serotype sample (serotype 17F). 1a. CPS probe intensities (normalised) 1b. Boxplot of probe intensities grouped by *cps* genes (only top 75 genes (out of 441 genes), with the highest median probe log intensity plotted for clarity.). Those *cps* genes which occur in serotype 17F are shaded black. 1c. *p*-values from t-tests of each of these 75 *cps* genes’ probes against all other probes. should go here.

Each serotype of *S. pneumoniae* contains a small subset of the 441 *cps* genes. The number of *cps* genes present in a serotype varies from 1 to 22, with an average of 13. There is considerable overlap in the gene complements of serotypes, with no serotype having a unique set of *cps* genes. In Figure 2b the *cps* genes of the serotype found in the sample being tested by this microarray are marked in black. It can be seen that some *cps* genes that do not occur in this serotype also have elevated intensities. This is due to cross-hybridization of probes. This occurs when it has not been possible to design probes entirely specific to a particular gene, so probes for one gene bind to some extent to the DNA from a different gene.

It is not uncommon for naso-pharyngeal samples to contain more than one serotype of *Streptococcus pneumoniae*. Figure 3 shows a boxplot of the CPS probe intensities from one microarray testing a sample that was found to contain three serotypes, 23F, 14 and 15C, in proportions 69%, 23% and 8%. Only the *cps* genes found in the three serotypes are shown in the figure for clarity. It can be seen that some of the *cps* genes are found in more than one of the serotypes contained in this sample, and one gene is found in all three.

**Figure 3.**
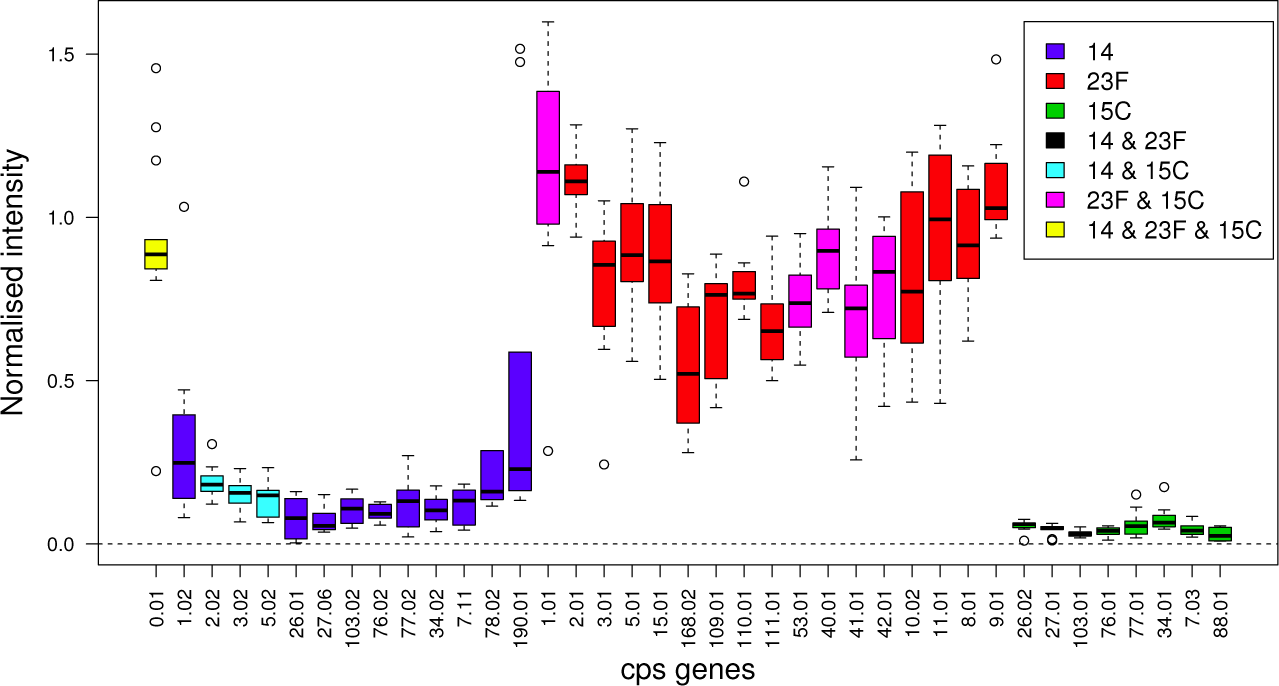
Array data from a typical sample containing a mixture of serotypes. The sample contains serotypes 23F, 14 and 15C in proportions 69%, 23% and 8%̤

Whilst the CPS probes are the primary measurements on the array for determining serotype content, eighteen sets of closely related serotypes have identical, or nearly identical, sets of *cps* genes, so cannot be distinguished by these probes alone. In order to differentiate between these closely related serotypes, the microarray contains extra probes, here referred to as STIDs. STID probes for a pair of serotypes come in pairs, with one probe for one serotype and a paired probe for the corresponding region of the genome of the second serotype. To test which of the two serotypes is present the difference in the fluorescent intensities of the paired probes is used. On average there will be around 65 pairs of STID probes for a pair of serotypes.

The arrays also suffer inevitably from different types of experimental noise. The fluorescent intensity signals can be affected by a variety of random factors which are difficult to control: from variation in the DNA extraction to variation in the binding of the DNA to its oligonucleotide probe due to divergent sequence. As mentioned above a probe’s intensity may be affected by cross-hybridisation; binding to DNA from a different gene. However in addition, the *Streptococcus pneumoniae* sample can be contaminated by DNA from the host and from other commensal or pathogenic organisms. This could also bind to some extent to a probe, affecting the probe’s measured intensity.

### Datasets

The dataset used in this work is an amalgamation of data from 60 different studies comprising more than four thousand samples. Every sample has initially been analysed by an empirical Bayesian Model and the result checked for errors by a single expert.

A sample may contain between 1 and 5 serotypes. Figure 4 shows a histogram of the frequency of occurrence in the dataset of samples containing different numbers of serotypes. The dataset was split into two, those arrays containing single serotypes and those arrays containing mixtures of serotypes. The single serotype arrays are used for training so were further filtered according to the quality-control information available. If the array had been flagged as suffering from low intensity, saturation, spatial problems or splashover it was removed. The dataset of samples containing mixtures of serotypes was not filtered by quality control information. This resulted in 3721 single serotype arrays and 926 mixture arrays.

**Figure 4.**
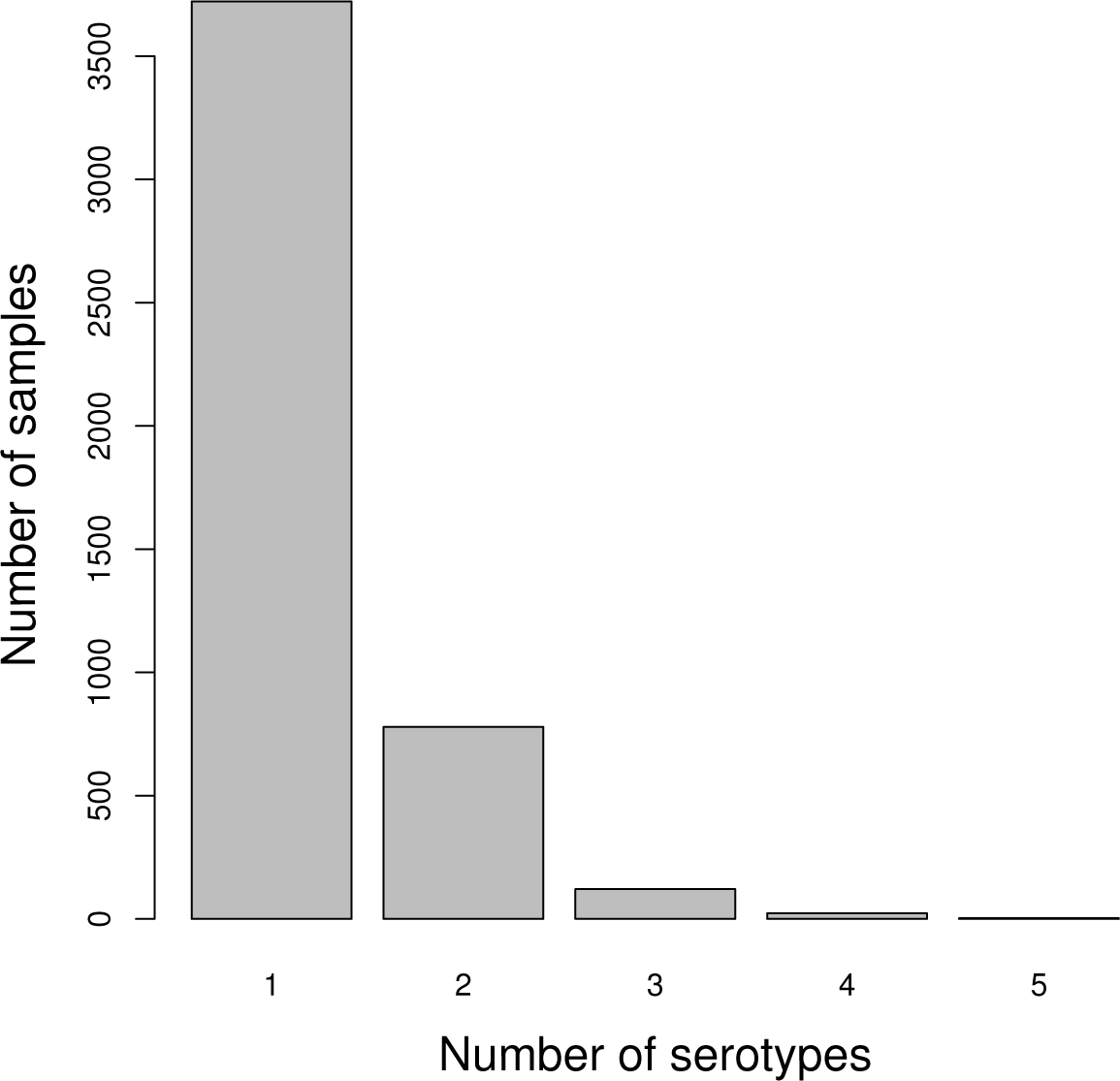
Number of samples containing different numbers of serotypes. Histogram showing the number of samples in the dataset containing different numbers of serotypes.

In the dataset of single serotype arrays the frequency of occurrence of serotypes is very variable, there are many examples of some serotypes, whilst very few of some others. The histogram in Figure 5 shows the number of single serotype arrays available for each serotype. Only 73 of the 91 known serotypes do feature in the single serotype array data, 18 rare serotypes have no examples at all. Hence we termed the overall dataset D.73. Given that the amount of training data can be critical for the performance of machine learning algorithms we started with a subset of serotypes that had at least 20 arrays in the single serotype dataset. This comprised a subset of 36 of the most frequent serotypes, and a total of 3486 single serotype arrays. We filtered the dataset of arrays containing mixtures of serotypes so that only these 36 serotypes featured in any sample, giving a total of 693 arrays. We termed the dataset of single serotype and mixture arrays that only included these 36 most common serotypes D.36. Details of D.73 and D.36, the number of arrays present and the number of serotypes featured in each, are tabulated in Table 1. The data used in this work is available from the authors on request.

**Figure 5.**
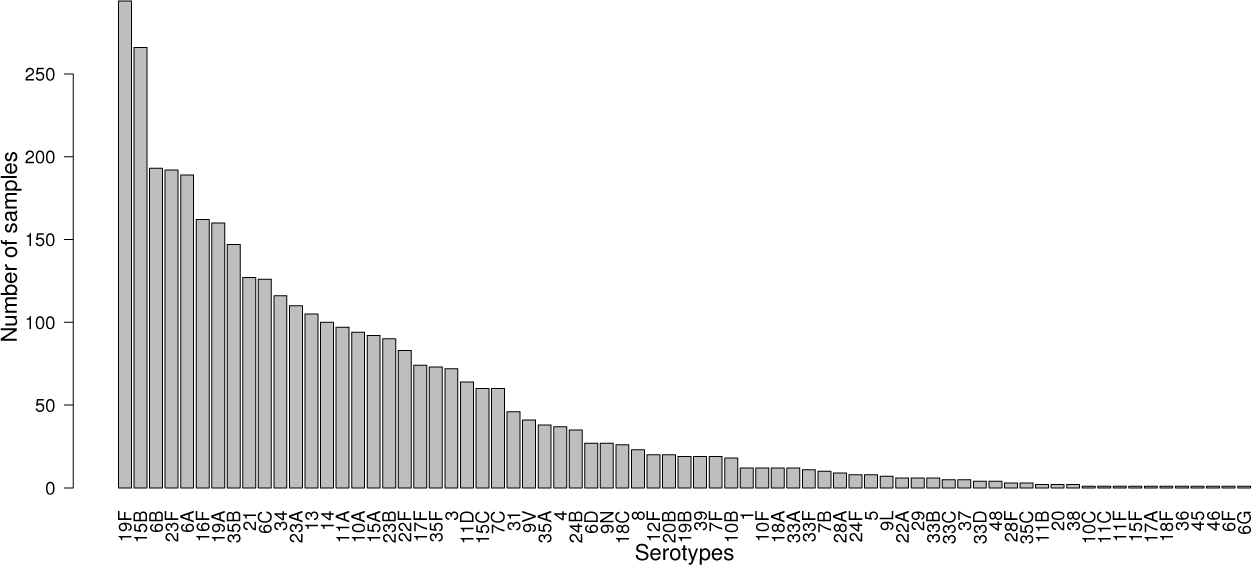
Number of single serotype arrays. Histogram showing the number of single serotype arrays in the D.73 dataset for each of the 73 serotypes.

**Table 1.** Details of datasets

| Name | Serotypes | Single arrays | Mixture arrays | Total arrays |
| --- | --- | --- | --- | --- |
| D.36 | 36 serotypes | 3486 | 693 | 4179 |
| D.73 | 73 serotypes | 3721 | 926 | 4647 |

### Pre-processing

For the Bayesian model no pre-processing was applied, since each array is analysed on its own without reference to other arrays. For the machine learning algorithms the arrays were first normalised. To normalise an array two reference levels were calculated; the median of the CPS probes on the array (*X*_*cps*_) and the median of the genome probes (*X*_*gen*_). All values *x* on the array were then normalised by *x*_*norm*_ = (*x−X*_*cps*_)/(*X*_*gen*_ −*X*_*cps*_). The normalised values were then logged. Initially the machine learning algorithms were applied to these CPS probe values directly, but the large number of probes resulted in relatively slow processing times. Therefore the CPS probe values were summarised to give one value for each of the 441 *cps* genes. Different summary methods were experimented with - mean, median, t-statistic and *p*-values. The *p*-values were found to work consistently well so these were adopted as the *cps* summary measure throughout the study. For each *cps* gene the *p*-values were calculated by testing whether the mean of the gene’s CPS probes was greater than the mean of all other CPS probes. The STID probes for a pair of serotypes are paired so the relevant measure is the difference in value between each probe pair. For each STID differentiated pair of serotypes there are approximately 65 such difference values, which were summarised by the *p*-value from testing whether their mean was significantly greater than zero. The R statistical environment [4] was used for all processing and analyses in this study.

### Empirical Bayesian Model

The Empirical Bayesian statistical model for calculating the probabilities of serotype combinations is described in detail in Newton et al [1]. To summarise we first set up likelihoods for gene binding depending on microarray log intensities for *cps* genes. Then, likelihoods of serotype combinations depending on gene binding and incorporating cross-hybridisation effects are derived. Further, likelihoods for serotypes depending on log intensities of STID probes are provided. Finally, all these likelihoods are put together to give a likelihood of serotype combinations depending on log intensities from CPS probes and STID probes. Combined with a prior on serotype combinations this allows us to infer a posterior probability for serotype combinations, apart from a normalising constant. Some of the hyperparameters of the model are estimated in an empirical Bayes fashion from the microarray data. Since there are exponentially many combinations of serotypes, we use a heuristic to limit the number of combinations to a subset of serotypes and serotype combinations with a potential for high probabilities.

## Machine learning

### Overview

In the machine learning approach to classification the algorithm is trained on one set of data, whose classes are known. The model learnt by the training step is then used to predict the classes of unknown data. In this application the data are the untransformed *cps* gene and STID *p*-values of the arrays and the classes are the serotypes on the arrays.

Both training and test data are needed to assess the performance of the machine learning algorithms. Therefore we performed 20-fold cross-validation on the single serotype array dataset; that is we divided the dataset into 20 equal parts, trained on 19 of the 20 combined and tested on the remaining part, repeating for each of the 20 parts. For the mixture arrays we trained on all the single serotype arrays and tested on the mixture arrays.

A sufficient amount of training data is required; enough training data to adequately represent the variability that the algorithm will encounter when tasked to classify test data. Whilst we had a considerable number of single serotype arrays to make training a single serotype classifier a feasible proposition this was not the case for mixture arrays. Being able to detect mixtures of serotypes in samples is an essential requirement for any serotyping method. Whilst the dataset D.36 (see Table 1) contained 693 mixture arrays this is only a small fraction of the number of possible combinations of 36 serotypes even if we restrict the maximum number of serotypes in a mixture to a realistic figure of between 2 and 5. In addition, for the serotype mixtures that we do have arrays for, there are then only a small number of the possible proportions of the mixtures represented. The solution we adopted is the two-step iterative method described below.

### Two-step method

In the two-step method we train the machine learning algorithm on a set of single serotype arrays (for testing mixture arrays this would be all the single serotype arrays, for testing single serotype arrays the 20-fold cross-validation scheme was used). Then, to test an unknown array, the first step is to use the classifier that has been generated by the training to predict the single serotypes on the unknown array. Then in the second step:

1. Choose the top *K* most probable predicted single serotypes.
2. Form all 2^*K*^ − 1 combinations (*K*_*c*_) of these K serotypes.
3. For each of these combinations:

- calculate the proportions of the constituent serotypes in the test array data from the medians of the serotypes’ *cps* genes’ probe values.
- Construct *M* artificial *mixture* arrays by combining randomly selected single serotype arrays, corresponding to the constituent serotypes, in the calculated proportions (setting *M* to 200 was sufficient in practice). The artificial mixture arrays were constructed by combining the single serotype arrays at the probe level. That is, in the artificial array a probe value is the weighted mean of the corresponding probe values in the single serotype arrays, weighted by the proportions calculated in point 3. The probe values are then summarised as *cps* gene *p*-values as described in the ‘Pre-processing’ section above.
4. Concatenate the *M* artificial arrays for the *K*_*c*_ combinations to create a new training data set.
5. Train a new classifier on these artificial mixture arrays.
6. Re-test the unknown array with the new classifier and choose the most probable class, giving the predicted serotype combination for the array.

Before choosing the top *K* most probable predicted serotypes (point 1 above) the list of predicted serotypes is edited for serotypes which are separated by STIDs; for pairs of serotypes that are separated by STIDs the serotype that has the lowest predicted probability is removed from the list.

In practice *K*, the number of most probable predicted serotypes (point 1 above), was chosen dynamically. Firstly the *cps* gene *p*-values where Bonferroni adjusted. Then for each serotype in the list of predicted serotypes, the percentage of that serotype’s *cps* genes that have adjusted *p*-values less than 0.05 was calculated. The serotypes with more that 50% of their *cps* genes significantly present were chosen for the second step. In practice *K* was rarely more than one more than the actual number of serotypes on the array. A limit of *K <*= 6 was set to prevent excessive processing times occurring.

### Filtering *cps* gene calls

The Machine learning algorithms in particular were found to be overly sensitive so a final filtering step can be used to remove False Positives from an algorithm’s call. The filtering step is based on the percentage of a serotype’s *cps* genes that are significantly present. For each array and each method the serotypes in the call from that method are examined in turn. The Bonferroni adjusted *p*-values of the *cps* genes are used to assess whether each gene is present or not. If a gene’s *p*-value is below a significance level of 0.05 the gene is deemed to be present. If a serotype has less than a pre-determined percentage of it unique *cps* genes present then the serotype is removed from the call. Note that only the *cps* genes that are *unique* to a serotype are considered. In a call containing several serotypes, in general some *cps* genes will be common to two or more serotypes, whilst some will be unique to each individual serotype; Figure 3 illustrates the point. In this filtering step it is the percentage of a serotype’s unique *cps* genes that are significantly present which is calculated.

This form of filtering was also used for a second role, that is, to divide the classified arrays into two categories with different confidence of prediction. For a sample and classification method, we inspected the serotype calls. For each serotype in the call we recorded the percentage of the serotype’s unique *cps* genes that were significantly present. If all the serotypes in the call had more than a predetermined percentage threshold of *cps* genes present, then that sample was placed in the high confidence group of samples.

### Evaluating results

The results for each method were compared to curated results to give error rates for each method, applied to each of the datasets. The error rates used were the percentage of arrays in the dataset which contained at least one False Positive serotype call or False Negative call or both. Credible intervals for the error rates were calculated using the 0.025 and 0.95 quantiles of the beta distribution. The curated results were based on the Bayesian Model results, which were used as a guide by the expert who inspected the arrays. Therefore there is the danger that the comparison will be biased in favour of the Bayesian Model over the Machine learning algorithms. We were however interested in ascertaining whether the Machine learning algorithms perform better than the Bayesian model so any bias will be acting to reduce rather than enhance the outcome we were looking for.

### Implementation

For the purpose of predicting sets of serotypes from *p*-value vectors we turned to two representative and popular Machine learning methods. The implementations of Random Forests and Gradient Boosting Machines contained in the machine learning platform H2O [5] were used, through the R [4] interface to the H2O platform provided by the R package ‘h2o’ [6].

The algorithms’ parameters were optimised using a grid search. Performance of different parameter combinations was assessed from the sum of two error rates. One was the error rate from the confusion matrix generated by the single serotype training data, that is, the performance of the algorithm as a single class classifier. Secondly the resulting classifier was applied to the mixture arrays. For each mixture array the list of the top six most probable predicted serotypes was examined to see if it contained all of the serotypes actually occurring in the sample. If all the actual serotypes were contained in the top six the test array was passed as not being an error, but if this was not the case the array was recorded as an error. The grid search minimised these two error rates combined. The parameter values used are shown in Table 2. The performance of the GBM as a single class classifier was, for this data, rather insensitive to parameter values, giving zero error rate using the default parameters. This performance was the same for any of the cross-validation subsets of the data as for the whole dataset. The parameters needed to be altered slightly from the default values in order to minimise the second error rate. The optimum parameter adjustments were the same if the classifier was trained on the whole dataset or on one of the cross-validation subsets.

**Table 2.**
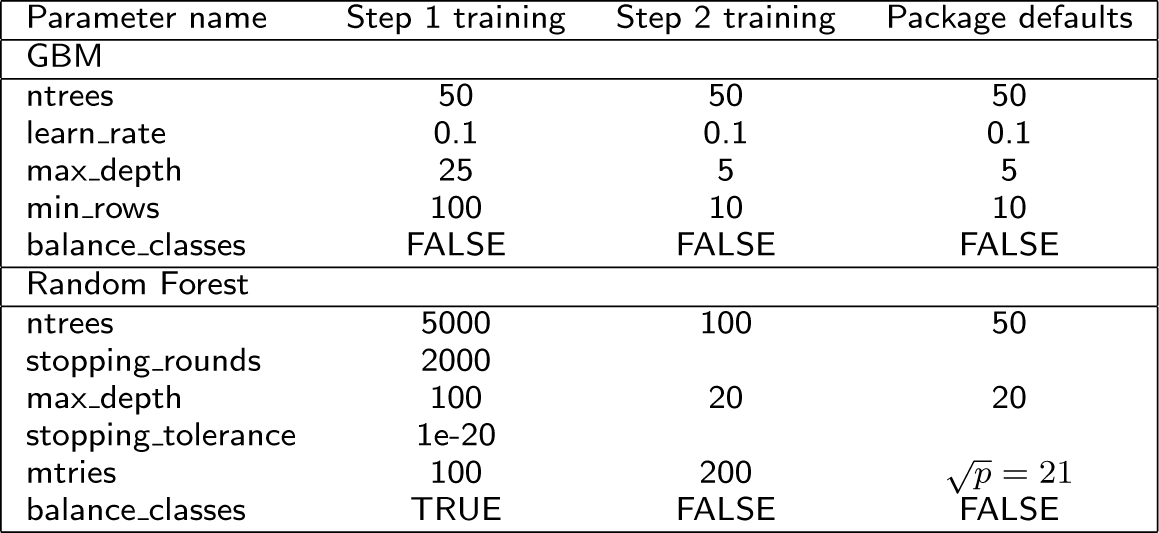
Machine Learning parameter values (*p* = **the number of input variables**)

| Parameter name | Step 1 training | Step 2 training | Package defaults |
| --- | --- | --- | --- |
| **GBM** |  |  |  |
| ntrees | 50 | 50 | 50 |
| learn_rate | 0.1 | 0.1 | 0.1 |
| max_depth | 25 | 5 | 5 |
| min_rows | 100 | 10 | 10 |
| balance_classes | FALSE | FALSE | FALSE |
| **Random Forest** |  |  |  |
| ntrees | 5000 | 100 | 50 |
| stopping_rounds | 2000 |  |  |
| max_depth | 100 | 20 | 20 |
| stopping_tolerance | 1e-20 |  |  |
| mtries | 100 | 200 | $\sqrt{p} = 21$ |
| balance_classes | TRUE | FALSE | FALSE |

For the Random Forest as a single serotype classifier the parameters needed to be changed from the default settings. In particular, balancing the classes was important for reducing the error rate. Perfect single serotype classification was not achieved but the error rate was around 0.1%. Again this did not vary whether applied to a cross-validation subset or to the whole dataset. Further adjustment of parameters was carried out to minimise the second error rate. The optimum parameter adjustments were the same if the classifier was trained on the whole dataset or on one of the cross-validation subsets.

The main speed constraint in the code is the fact that multiple t-tests are required. 441 tests are needed for each of the *K*_*c*_ x 200 arrays, where *K*_*c*_ could be as high as 63. The time taken for these was minimised using the parSapply routine from the R package doParallel [7], to distribute the 441 calculations between computer cores. The routine rowttests from the R package genefilter [8], was used for the t-tests. This routine is implemented in C for speed, and was edited to perform a t-test where the two groups being tested have unequal variance. On average one array takes less than one minute to analyse on a 4-core 2GHz machine. The R code used in this paper is available from the authors on request.

### Artificial single serotype training arrays

The data currently available contains no examples of 18 particularly rare serotypes and few (less than 20) examples of 37 other serotypes. We investigated the possibility of generating entirely artificial *single* serotype arrays to be used for training a classifier for the rare serotypes. For each of the single serotype arrays, we recorded the CPS probe values for the *cps* genes known to occur in that serotype, and the CPS probe values for all other *cps* genes on the array. We concatenated these two sets of probe values for all the arrays to produce two distributions - a distribution of probes for genes present (the alternative distribution) and a distribution of probes for genes absent (the null distribution). To create an artificial array for a particular serotype we took the known *cps* gene profile for the serotype in question and sampled from the alternative distribution to generate CPS probe values for those genes, and sampled from the null distribution to generate CPS probe values for all other genes. In this way we created 200 single serotype arrays for each serotype. In order to be able to validate the performance we explored the idea using the D.36 dataset. We used this artificial single serotype dataset to train a classifier and predict the D.36 arrays using the two-step process described above and compared the results with those obtained using a classifier trained on the actual D.36 single serotype arrays.

### Results

#### Comparing methods

The error rates for the three methods, the Bayesian Model, GBM and Random Forest, are shown in Table 3 for the datasets D.36 and D.73 (see Table 1 for dataset details). For the D.36 dataset the GBM performs much better than the Bayesian Model for samples containing mixtures of serotypes, with the Random Forest performance intermediate between the two. For single serotype samples however the performance of all three methods is approximately the same, noting the credible intervals. The error rate for the whole D.36 dataset, single and mixture arrays combined, is slightly better for the GBM and roughly the same for the Bayesian Model and Random Forest; since there is a preponderance of single serotype arrays in the dataset.

**Table 3.** Percentage error rates for the Bayesian Model (BM), GBM and Random Forest (RF) for the datasets defined in Table 2; for samples containing mixtures, samples containing single serotypes and all samples combined. Figures in brackets show the 95% credible interval.

| Dataset |  | BM |  | GBM |  | RF |  |
| --- | --- | --- | --- | --- | --- | --- | --- |
| D.36 | Mixtures | 18 | (15.2 - 21.0) | 5.6 | (4.0 - 7.4) | 10.0 | (7.9 - 12.3) |
|  | Singles | 8.1 | (7.2 - 9.0) | 8.3 | (7.4 - 9.2) | 9.8 | (8.8 - 10.8) |
|  | Combined | 9.7 | (8.8 - 10.6) | 7.9 | (7.1 - 8.7) | 9.8 | (8.0 - 9.7) |
| D.73 | Mixtures | 21 | (18.4 - 23.7) | 19 | (16.5 - 21.6) | 21 | (18.4 - 23.7) |
|  | Singles | 8.2 | (7.3 - 9.1) | 22.0 | (20.7 - 23.3) | 20.0 | (18.7 - 21.3) |
|  | Combined | 10.8 | (9.9 - 11.7) | 21.4 | (20.2 - 22.6) | 20.2 | (19.1 - 21.4) |

For the D.73 dataset, which includes serotypes with little training data, the three methods give roughly the same results for mixture arrays. The Bayesian Model, unaffected by lack of training data, has the same error rate as for D.36, but the machine learning error rates are increased greatly, reflecting insufficient training data for many serotypes in the dataset. For single serotype arrays, again the Bayesian Model has a similar error rate to D.36 but the two machine learning methods error rates are much worse. This suggests that at least 20 training examples are needed for successful machine learning of serotyping data.

However, as described in the methods section, a further processing step can be applied to the results in order to reduce the number of False Positives; filtering based on the percentage of a serotype’s unique *cps* genes that are significantly (*p*-values *<* 0.05) present. For each sample and for each method, the serotypes in the method’s call are examined to see what percentage of their *cps* genes are significantly present. If fewer *cps* genes are significantly present than a set percentage threshold then this serotype is removed from the method’s call.

Figure 6 shows the effect of this filtering using thresholds from 0% to 100% on the D.36 dataset. It can be seen that this filtering can improve the error rates of all three methods, as falsely positive serotypes are removed from the calls, without a concomitant increase in false negatives. The results from selecting a threshold of 50% are shown in Table 4. For mixtures samples in the D.36 dataset the error rates are not greatly changed, but for the single serotype samples the error rates are reduced, in particular for the GBM with an error rate now of only 1.5%. For the D.73 dataset, filtering gives improvements for all three methods for mixture and single serotype samples, with large improvements for the machine learning algorithms on the single serotype arrays, since the errors seen in the unfiltered results of Table 3 are mainly false positives that are removed by the filtering.

**Figure 6.**
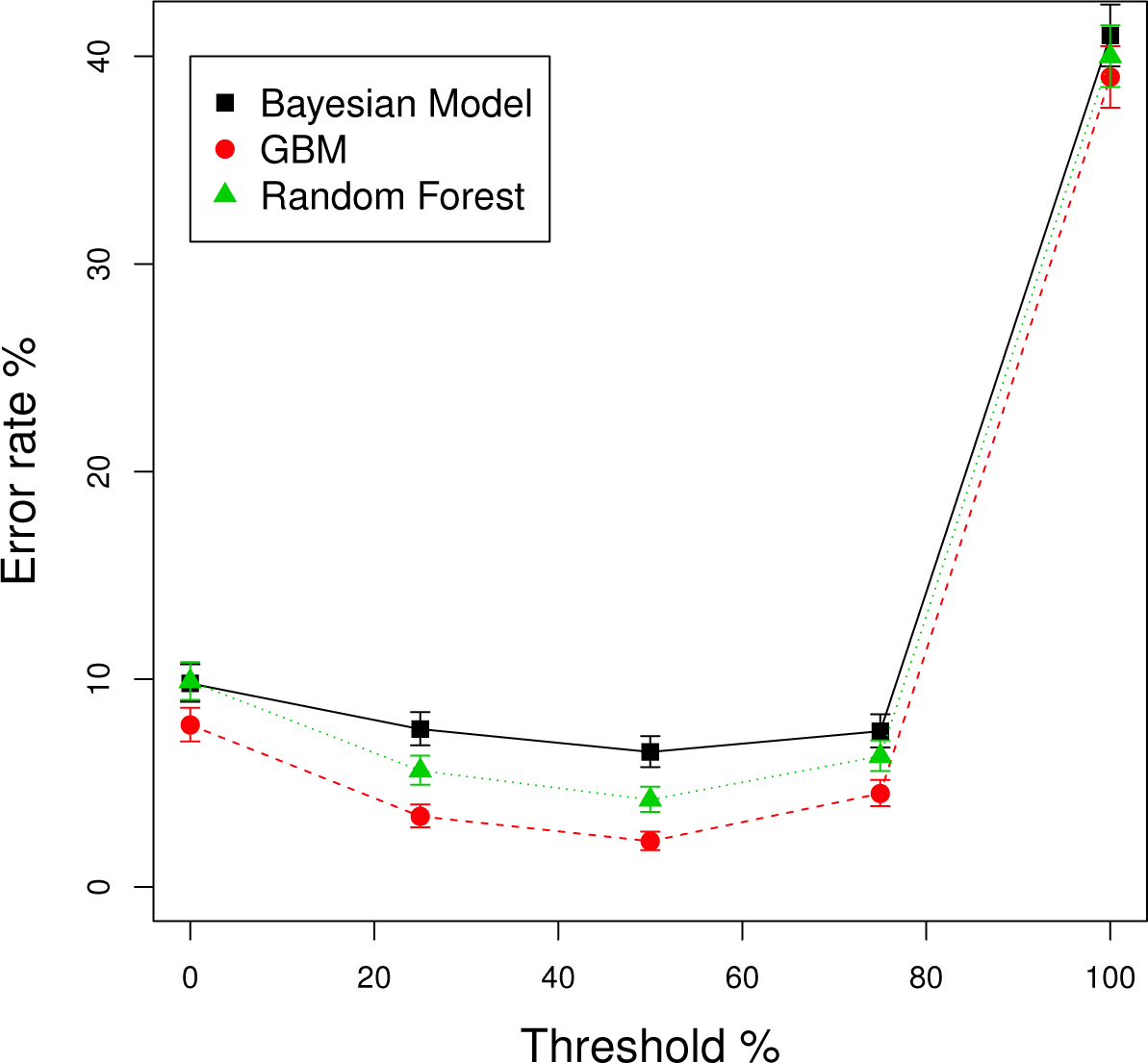
Error rates after filtering calls based on the percentage of cps genes present. The effect on error rates of filtering serotype calls based on the p ercentage of a serotype’s *cps* genes that are significantly present. Different thresholds of percentage of genes present were used and plotted along the x-axis. Error bars are the 95% credible intervals. D.36 dataset.

**Table 4.** Percentage error rates for the Bayesian Model (BM), GBM and Random Forest (RF) after filtering calls on the percentage of a serotype’s *cps* genes significantly present, using a 50% threshold; for samples containing mixtures, samples containing single serotypes and all samples combined. Figures in brackets show 95% credible interval.

| Dataset |  | BM |  | GBM |  | RF |  |
| --- | --- | --- | --- | --- | --- | --- | --- |
| D.36 | Mixtures | 15 | (12.4-17.8) | 5.9 | (4.3-7.8) | 9.1 | (7.1-11.3) |
|  | Singles | 4.8 | (4.1-5.5) | 1.5 | (1.1-1.9) | 3.3 | (2.7-3.9) |
|  | Combined | 6.5 | (5.8-7.3) | 2.2 | (1.8-2.7) | 4.3 | (3.7-4.9) |
| D.73 | Mixtures | 19 | (16.5-21.6) | 12 | (10.0-14.2) | 16 | (13.7-18.4) |
|  | Singles | 5.6 | (4.9-6.4) | 5.0 | (4.3-5.7) | 7.1 | (6.3-7.9) |
|  | Combined | 8.3 | (7.5-9.1) | 6.4 | (5.7-7.1) | 8.9 | (8.1-9.7) |

#### Artificial single serotype training data

We investigated the possibility of generating entirely artificial *single* serotype arrays to be used for training a classifier for the rare serotypes. In order to be able to validate the approach we used the D.36 dataset. For simplicity we removed all arrays containing serotypes with STIDS. This left 28 serotypes and 3486 single serotype arrays and 339 mixture arrays. The results are shown in Table 5, with the results from the Bayesian model and from the GBM trained on actual single serotype arrays included for comparison. Overall the approach does not perform well with a 12.8% error rate for all arrays compared to 6.1% for the Bayesian Model (with filtering included). For predicting mixture arrays the artificial training data approach performs particularly badly (error rate of 45%). For predicting single serotype arrays, provided filtering is used, the approach gives a slightly worse performance (8.4%) than the Bayesian model (5.0%).

**Table 5.**
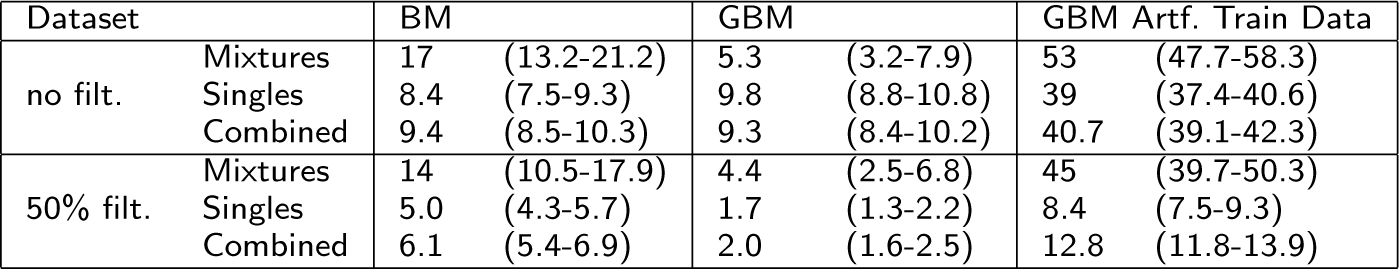
Results for the artificial single serotype training data. Percentage error rates for the Bayesian Model (BM), GBM using actual single serotype arrays for training (GBM) and GBM using artificial single serotype arrays for training (GBM Artf. Train Data). 50% filtering refers to filtering calls on the percentage of a serotype’s *cps* genes significantly present using a 50% threshold. For samples containing mixtures, samples containing single serotypes and all samples combined.

#### A practical implementation

The GBM gives a lower error rate than the Bayesian Model, however a paucity of training data for some serotypes in the D.73 dataset does increase the GBM error rate. In addition 18 known serotypes which could be encountered in clinical samples currently lack any training data at all. An attempt to create artificial training data was unsuccessful, so until sufficient actual training data is available for all serotypes the Bayesian model will have to be used to analyse the arrays. The Bayesian model is unaffected by lack of training data, giving similar error rates for both the D.73 and the D.36 dataset.

We investigated whether the GBM could be used to divide samples into two categories; a group of samples which we have high confidence in the class assignments (group C) and a second group of samples whose class assignments are less confident (group NC). There will clearly be a trade off, lowering the expected error rate that would be acceptable in group C, will increase the number of samples allocated to group NC.

We investigated two different methods for dividing the dataset into the two groups. Firstly, for each sample, if the serotype calls from the Bayesian model and from the GBM were identical then the sample was placed in group C, that is, the high confidence group. If the serotype calls differed then the sample was placed in group NC. For the D.36 dataset we found group C contained 84% of the dataset with an error rate of just 0.37%. For the D.73 dataset group C contained 72% of the dataset with an error rate of 0.69%. These results may be biased since they include the Bayesian model results and, as explained in the Methods section, the curated results used these as a guide. We therefore tried a second method for creating two groups that does not use the Bayesian model results.

The second method for creating two groups used a threshold on the percentage of a serotype’s *cps* genes significantly present. For each sample and each classification method separately, we inspected the serotype calls. For each serotype in the call we recorded the percentage of the serotype’s unique *cps* genes that were significantly present (that is, having a Bonferroni adjusted *p*-value less than 0.05). If *all* the serotypes in the call had more than a predetermined percentage threshold of *cps* genes present, then that sample was placed in group C, the high confidence group. If *any* of the serotypes had less than this percentage of genes present then the sample was placed in group NC. Percentage thresholds from 65% to 100% were tested. The percentage of samples placed in group C and the error rate in the group were recorded. The results for the GBM on the dataset D.36 are shown in Figure 7. We found that with a threshold of 90% the error rate in group C was just 0.31% and it contained 78% of the dataset. Reducing the percentage threshold increases the number of samples in group C, but also increases the error rate in the group. Increasing the threshold above 90% does not reduce the error rate further, but does reduce the number of samples in the group. Applying the same method to the Bayesian model placed 80% of the samples in group C with an error rate of 3.7%, again with a percentage filtering threshold of 90%.

**Figure 7.**
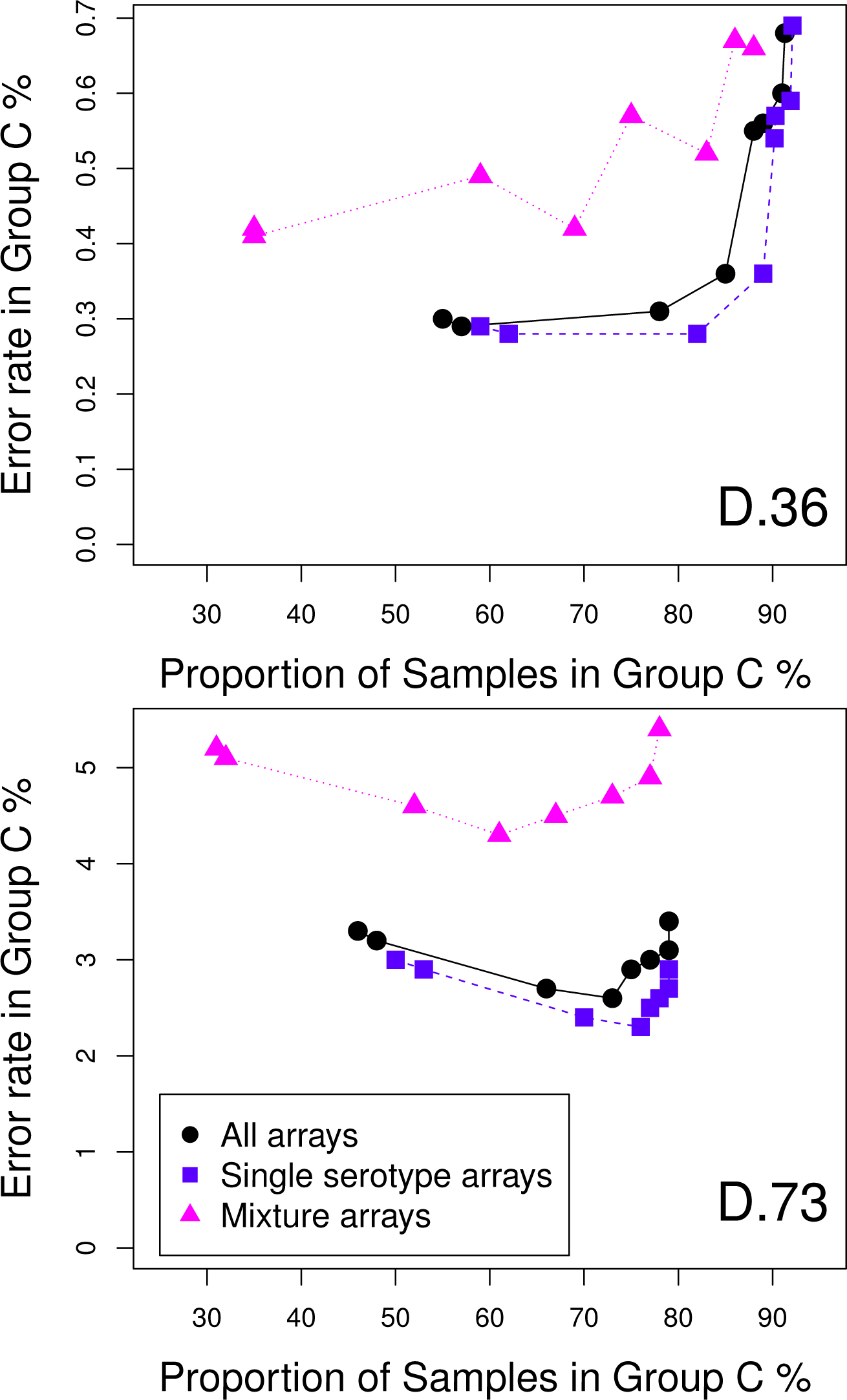
Error rate in the high confidence group of samples against the percentage of the dataset in that group. The GBM results and a threshold on the percentage of a serotype’ s *cps* genes significantly present were combin ed in order to divid e the dataset into two groups of sampl es, a high confidence group of sampl es (G roup C) and others (G roup NC) Th e graph plots the error rate in G roup C against th e percentage of the dataset in that group, as the threshold is varied For datasets D 36 and D 73

The results for the D.73 dataset are also shown in Figure 7. With a 90% threshold the error rate in group C is now 2.6% and contains 70% of the dataset’s samples. For the Bayesian model the values are 4.7% and 20%.

Finally we investigated combining the two different methods to define the two groups. We first divided the dataset based on the agreement between the Bayesian model and the GBM. We then examined the samples in group C using the second method for defining groups. Samples that had any called serotypes with a percentage of *cps* genes lower than a predetermined threshold were moved from group C to group NC. The results for the D.36 dataset are shown in Figure 8. Using a threshold of 65% gives a 0.17% error rate (8 errors) in group C which contains 83% of the dataset. With a threshold of 95% the error rate is 0.046% (1 error) with 52% of the samples in group C. The results for the D.73 dataset are also shown in Figure 8; with a threshold of 65% the error rate in group C is 0.49% (16 errors) which contains 70% of samples, with a threshold of 95% the error rate in group C is 0.25% (5 errors) with 42% of samples.

**Figure 8.**
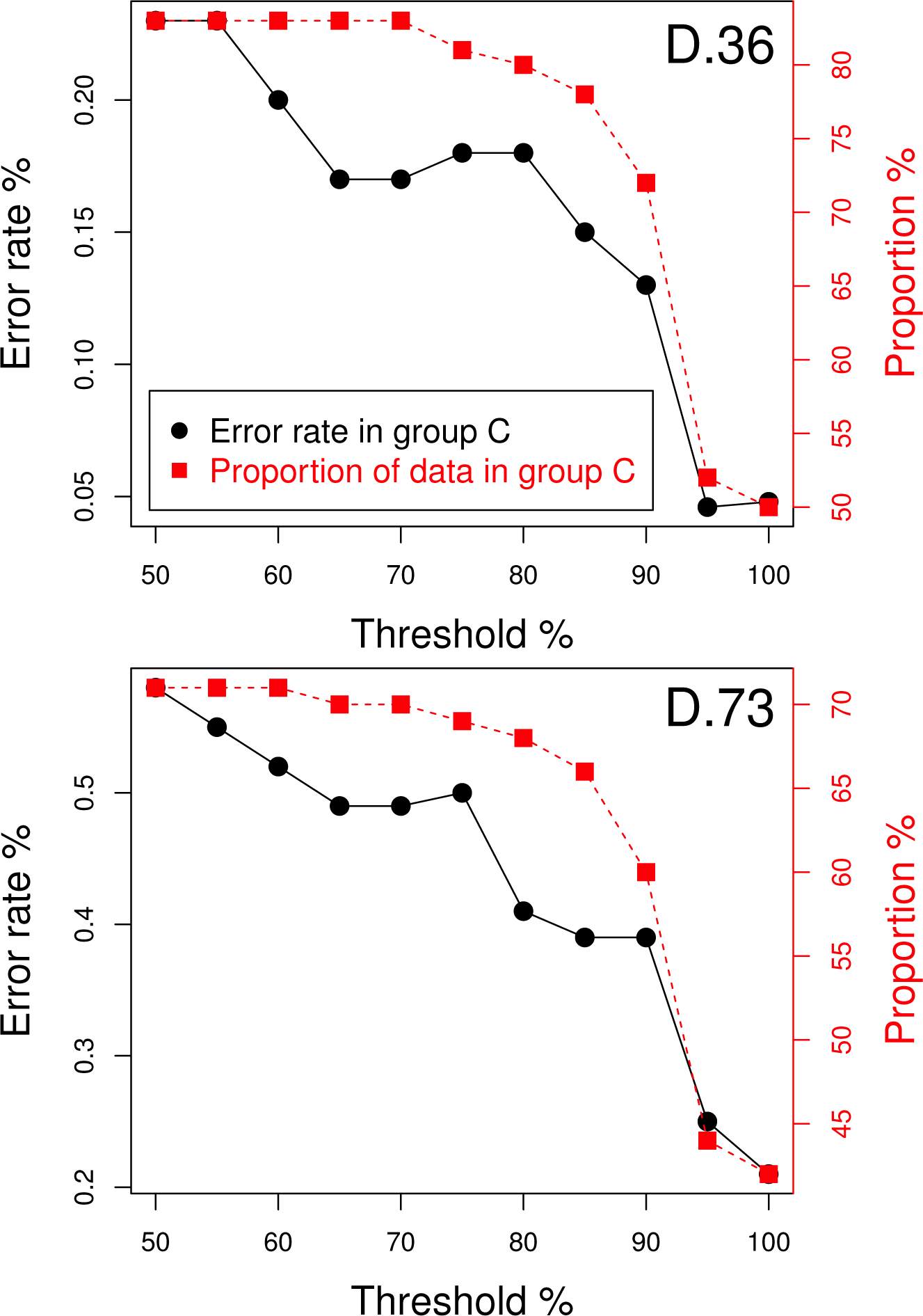
Results of combining two methods to divide the dataset into two groups of samples. Combining two methods to divide the dataset into two groups of samples, a high confidence group of samples (Group C) and others (Group NC), on e method being th e agreement betwee n the Bayesian model and the GBM, the other method being a threshold on the percentage of a serotyp e’s *cps* genes significantly present Graph plots th e error rate in Group Con th e l efty-axis and the proportion of the dataset in G roup C on the rig ht y-axis, against the threshold used in the second method. For datasets 0.36 and D. 73.

#### Variable importance

Training on the single serotype arrays gives the relative importance of the variables (*cps* genes) in classifying each serotype. The H2O [5] GBM algorithm calculates this for each variable based on whether that variable was selected during splitting in the tree building process and how much the squared error improved as a result [9].

Examining the variable importance can reveal biological insights into the *cps* structure of serotypes. For example serotypes 6A and 6B have identical *cps* gene profiles so are distinguished in the Bayesian model of the data by a STID. We found however that in more than 80% of cases it was possible to correctly classify 6A and 6B single serotype arrays without including the STID probes in the training data. So there are *cps* gene differences between the two closely related serotypes and examining the variable importances can reveal the nature of those differences.

We took all single serotype arrays that contained serotypes 6A and 6B, comprising 189 6A arrays and 192 6B arrays. We randomly sampled 25% of the arrays to form a test set and trained on the remainder. Only the *cps* gene *p*-values were used, not the STID *p*-values. After training and prediction we recorded the number of True Positive predictions. We performed a grid search of the training parameters to maximise the True Positive rate. The 25% sampling was performed 5 times for each grid point of parameters and the average True Positive rate recorded. Setting the number of trees to 50, maximum depth to 200 and minimum rows to 50 maximised the True Positive rate, giving 88% for 6A and 82% for 6B. Using these parameter values we ran the training and testing, with 25% sampling, 100 times and recorded the variable importances of the 441 *cps* genes each time. Figure 9 shows a boxplot of the 100 relative importance values for each *cps* genes, ordered by mean variable importance, with only the top 25 genes with the highest mean variable importance shown for clarity. The figure indicates which *cps* genes the GBM has learnt are important in distinguishing the two serotypes. One gene, 81.02, is consistently essential for distinguishing between the two serotypes and approximately six more have some importance.

**Figure 9.**
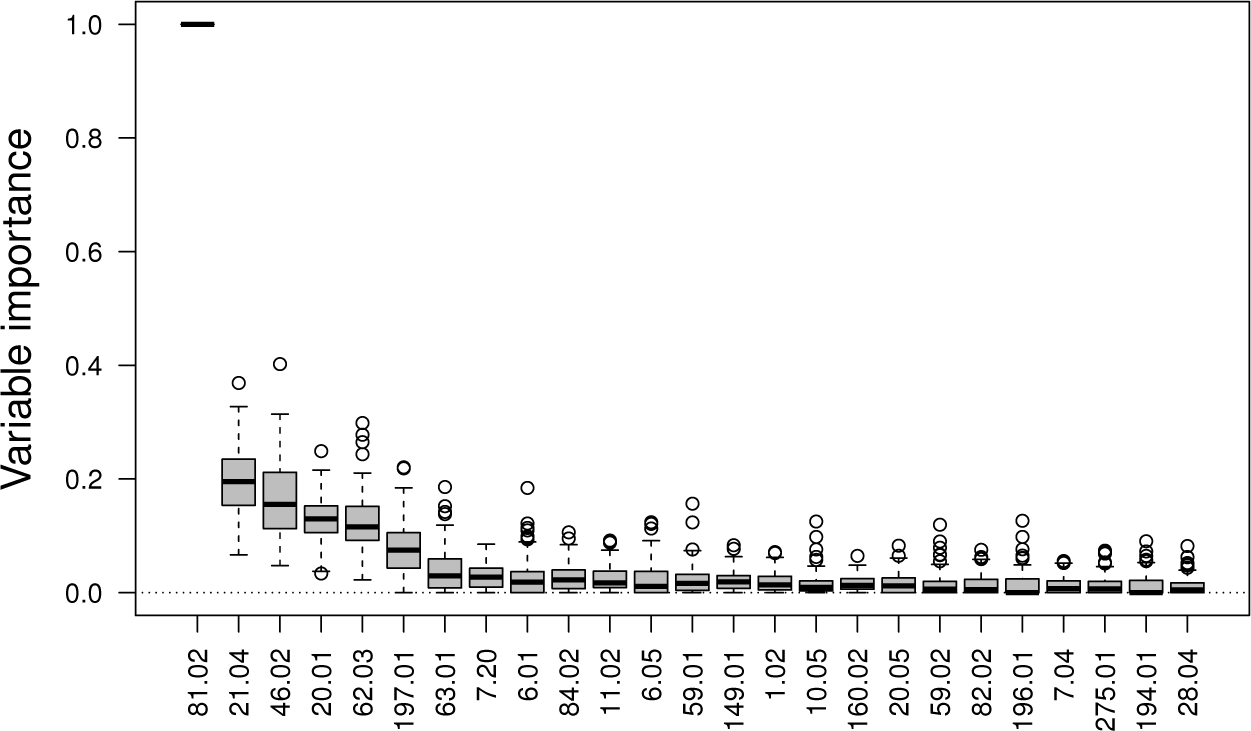
Variable (cps gene) importance for distinguishing serotypes 6A and 68. The top 25, out of 441, *cps* genes shown for clarity, ordered by their mean importance evaluated from 100 cross-validation samplings.

Investigating gene 81.02 in more detail. Figure 10 shows the mean, standard deviation and p-values for gene 81.02 from 189 arrays containing serotype 6A (red dots) and 193 arrays containing serotype 6B (black dots). It can be seen that gene 81.02 is significantly present in both 6A and 6B arrays, with all p-values less than 0.05 (horizontal line). However the gene’s probes have a noticeably lower median intensity and higher standard deviation in most 6B arrays compared to the values in 6A arrays, suggesting that there is some sequence divergence in gene 81.02 between serotypes 6A and 6B. This is an example of how machine learning algorithms easily pick up on subtle differences that humans may miss.

**Figure 10.**
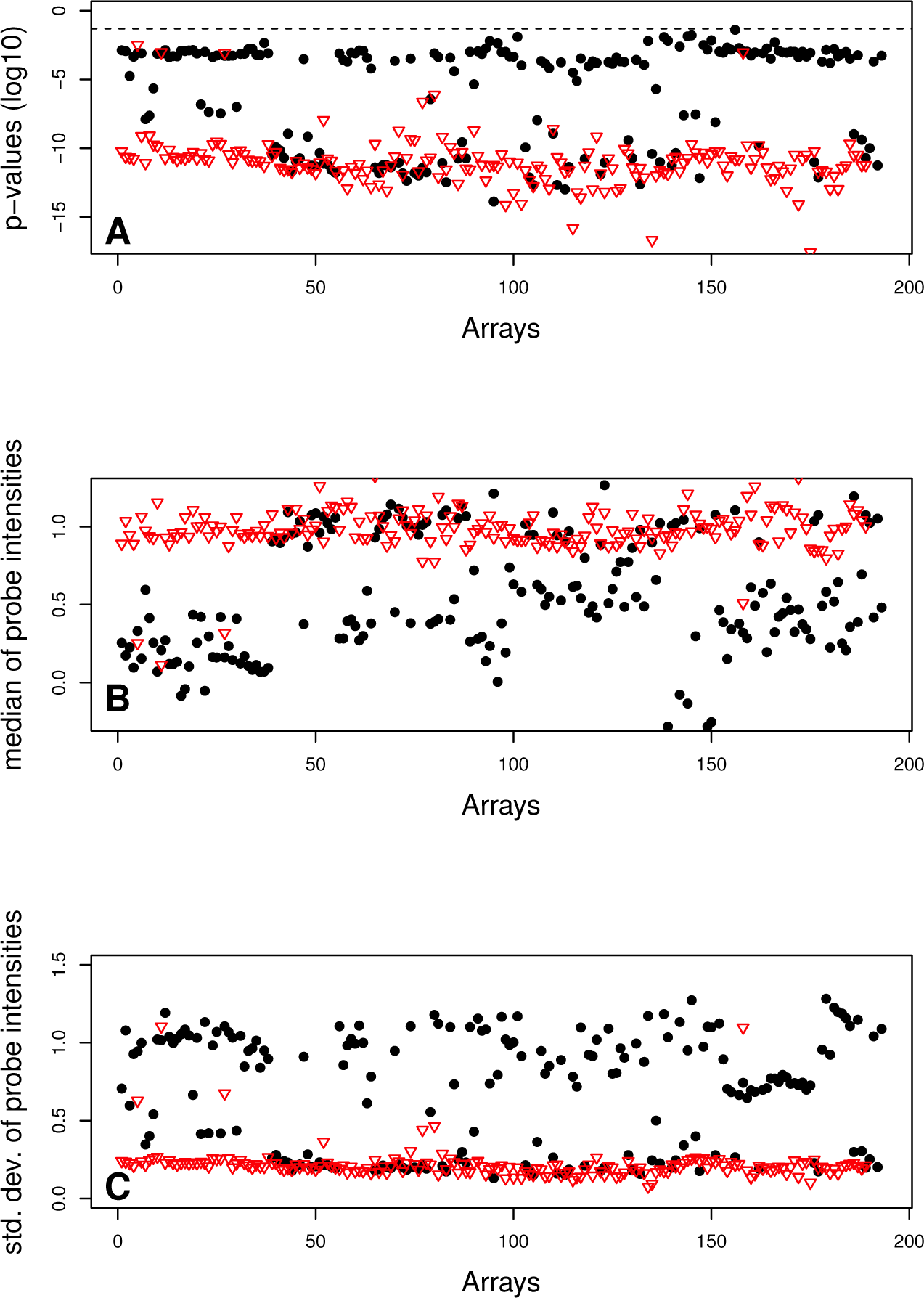
Summary values for gene 81.02 probes from arrays containing serotype 6A and 68. Summary values for gene 81.02 probes from 189 arrays containing serotype 6A (red dots) and 193 arrays containing serotype 6B (black dots) A p-values of probes (dotted lin e = log10(0 05)) B Median of probe intensities. C. Standard deviations of probe intensities.

### Discussion

A machine learning approach using a Gradient Boosting Machine can give a better error rate classifying serotyping data than even a carefully crafted Bayesian model. A lack of training examples containing mixtures of serotypes can be overcome with an iterative two-step approach that creates artificial mixture arrays from single serotype arrays. Sufficient single serotype training data is important, but even with relatively few training examples machine learning still gives a better performance than the Bayesian model provided an extra filtering step is included to remove False Positives. Of the two machine learning algorithms tested the GBM performed better than Random Forests.

The Bayesian model relies on known prior information such as the *cps* gene profiles of the serotypes. The strength of the machine learning approach is that there may be features in the data important for accurate classification which are currently unknown but which machine learning can identify automatically. And with sufficient training examples machine learning can cope with biological and experimental variability in features important for classification. The errors that do occur in the results from the GBM algorithm may be due to biological variability occurring in the erroneous samples, variability that the algorithm has not been sufficiently trained to detect.

Because training data is not available for all serotypes, in practice the Bayesian model will still need to be used to analyse the arrays, but we have shown that combining the results with those from the GBM can be used to assign arrays to a high confidence group with a very low error rate. The added advantage of a machine learning approach is that as further data becomes available the classifier can be updated with a view to increasing the prediction accuracy and increasing the proportion of arrays assigned to the high confidence group.

The bacterium is constantly evolving so it is likely that in the future the machine learning algorithm will encounter further serotypes which it will not have been trained to recognise. So probably an approach using the Bayesian model in combination with a GBM may never be entirely redundant. The advantage of the Bayesian model is that the *cps* gene profile of a new serotype can be added straight away to the information supplied to the Bayesian model, whereas the GBM will require training examples of the new serotype first.

When no training data is available we found the Bayesian model is better than a machine learning approach that uses artificial single serotype training arrays. Lacking probe data for genes unique to the missing 18 serotypes meant that probe specific distributions were not possible. The approach also ignored cross-hybridisation effects. In future work we will investigate whether it is possible to construct better artificial training data using the information from a BLAST search of all the array probes against the sequences of the serotypes’ *cps* genes.

### Conclusions

The study illustrates the strength of a machine learning approach for classifying genomic data with multiple variables. The algorithm learns which variables are important for classification, unlike in a modelling approach where the user has to make this choice. It also demonstrates that when suitable training data is lacking steps can be taken to create artificial training data. The general conclusion of the study however is that in practical classification problems the best approach may not be choosing between either a Bayesian modelling or a machine learning solution but using a combination of the two methods. The study also illustrates how the relative variable importance from a machine learning classification can be used as an efficient research tool to investigate differences in the objects to be classified, in this case the differences in the genomic compositions of the serotypes.

### Declarations

Ethics approval and consent to participate

Not applicable.

Consent for publication

Not applicable.

### Availability of data and material

The datasets used and/or analysed during the current study are available from the corresponding author on request.

## Authors’ contributions

The authors contributed equally to the work and read and approved the final manuscript.

## Acknowledgements

Not applicable.

